# Muller’s ratchet as a mechanism of frailty and multimorbidity

**DOI:** 10.1101/439877

**Authors:** Diddahally R. Govindaraju, Hideki Innan

**Affiliations:** Department of Human Evolutionary Biology, Harvard University, Cambridge, MA 02138; The Institute of Aging Research, Albert Einstein College of Medicine, Bronx, NY 1046, and; Graduate University for Advanced Studies, Hayama, Kanagawa, 240-0193, Japan

**Keywords:** Senescence, mutation accumulation, asexual reproduction, somatic cell-lineages, organs and organ systems, Muller’s ratchet, fitness decay, frailty and multimorbidity

## Abstract

Mutation accumulation has been proposed as a cause of senescence. In this process, both constitutional and recurrent mutations accumulate gradually and differentially among differentiating cells, tissues and organs, in relation to stage and age, analogous to Muller’s ratchet in asexually reproducing organisms. Latent and cascading deleterious effects of mutations might initiate steady “accumulation of deficits” in cells, leading to cellular senescence, and functional decline of tissues and organs, and ultimately manifest as frailties and disease. We investigated a few of these aspects in cell populations through modeling and simulation using the Moran birth-death process, under varied scenarios of mutation accumulation. Our results agree with the principle of Muller’s ratchet. The ratchet speed in a given tissue depends on the population size of cells, mutation rate, and selection coefficient. Additionally, deleterious mutations seem to rapidly accumulate particularly early in the life-course, during which the rate of cell division is high, thereby exerting a greater effect on cellular senescence. The speed of the ratchet afterward varies greatly between cells nested in tissues and tissues within organs due to heterogeneity in the life span and turnover rate of specific cell types. Importantly, the ratchet accelerates with age, resulting in a synergistic fitness decay in cell populations. We extend Fisher’s average excess concept and rank order scale to interpret differential phenotypic effects of mutation load in a given tissue. We conclude that classical evolutionary genetic models could explain partially, the origins of frailty, subclinical conditions, morbidity and health consequences of senescence.

**Significance:** Frailty is defined as physiological and functional decline of organs and organ systems, due to deficit accumulation from stochastic damages within the organism with advanced age. Equivalently, with age, both constitutional and somatic mutations accumulate gradually and differentially among cells, cell lineages, tissues, and organs. Since most mutations are deleterious, accumulation of random and recurrent mutations could create a “load,” on the genome and contextually express in the epigenome and phenotype spaces. Here we extend Muller’s ratchet principle to explain frailty and multi-morbidity using the Moran model and simulations. Our results agree with the Muller’s ratchet principle. We emphasize the need for considering cumulative effects of the entire spectrum of mutations for explaining the origin of frailty, sub-clinical conditions, and morbidity.

---

“Different organs of the body have their individual liabilities, as also same organ in different individuals … idiosyncrasy whether of individual subjects or of individual organs, is an important factor…” (1).

“If one organ or group, for any accidental reason begins to function abnormally and finally breaks down, the balance of the whole is upset and death eventually follows” (2).

**S**enescence is a time-dependent biological process involving irreversible arrest of cell proliferation, altered metabolic activity, functional deterioration of tissues, organs and organ systems of individuals and populations (3, 4). From a developmental genetics viewpoint, senescence or the biological aging process may be initiated shortly after the formation of the zygote (i.e., at the first mitotic division) and progress at variable pace into older ages, culminating in death (5–8). Organisms are the ultimate products of integrated hierarchical organization and development as well as mutually influencing modular systems ranging from molecules to organs and organ system (9, 10). Accordingly, the deteriorative processes associated with senescence could arise at any of these levels and potentially influence only a few or all levels of ontological hierarchies: cells, organs, organ systems and individuals at different rates (8, 11, 12). Ultimately, the cumulative effect of such deteriorative processes may lead to a functional decline of the correlated and interacting components of the biological system as well as affect viability (longevity) and reproduction of individuals - collectively, the Darwinian fitness.

## Senescence as an intrinsic property of developmental process

In brief, developmental processes are initiated by the fusion of male and female haploid gametes to form a unicellular and omnipotent diploid cell - the zygote. Following a series of mitotic divisions, zygote differentiates into three layers of pluripotent stem cells- endoderm, mesoderm, and ectoderm (13). Stem cells in these layers further undergo a succession of divisions, transformations and differentiation and get assorted spatially into quasi-independent, specialized, yet integrated hierarchical modules of tissues, organs and organ systems within the organism (14, 15) (**Figure 1**). As modular traits, they show ontogenetic and functional integration and interaction as well as influence each other across all levels of the genotype - epigenetic - phenotype (G-E-P) space (9, 10, 16–18). Fully differentiated and developed organs and organisms not only exhibit individual variation for specific metric traits, but each of these traits also generally follow a Gaussian distribution at population levels, a feature typical to quantitative traits. For instance, measures of kidney, heart, liver, brain, and lungs vary among individuals (19, 20). Thus, the individual may be viewed as spatially organized and interacting system of polygenic traits/organs. Clearly, each of these traits, and their components are subjected to age related changes, just as organisms do. Hence, senescence as an intrinsic aspect of developmental process would influence the G-E-P spaces of all organs, organs systems and ultimately, the individual.

**Figure 1:**
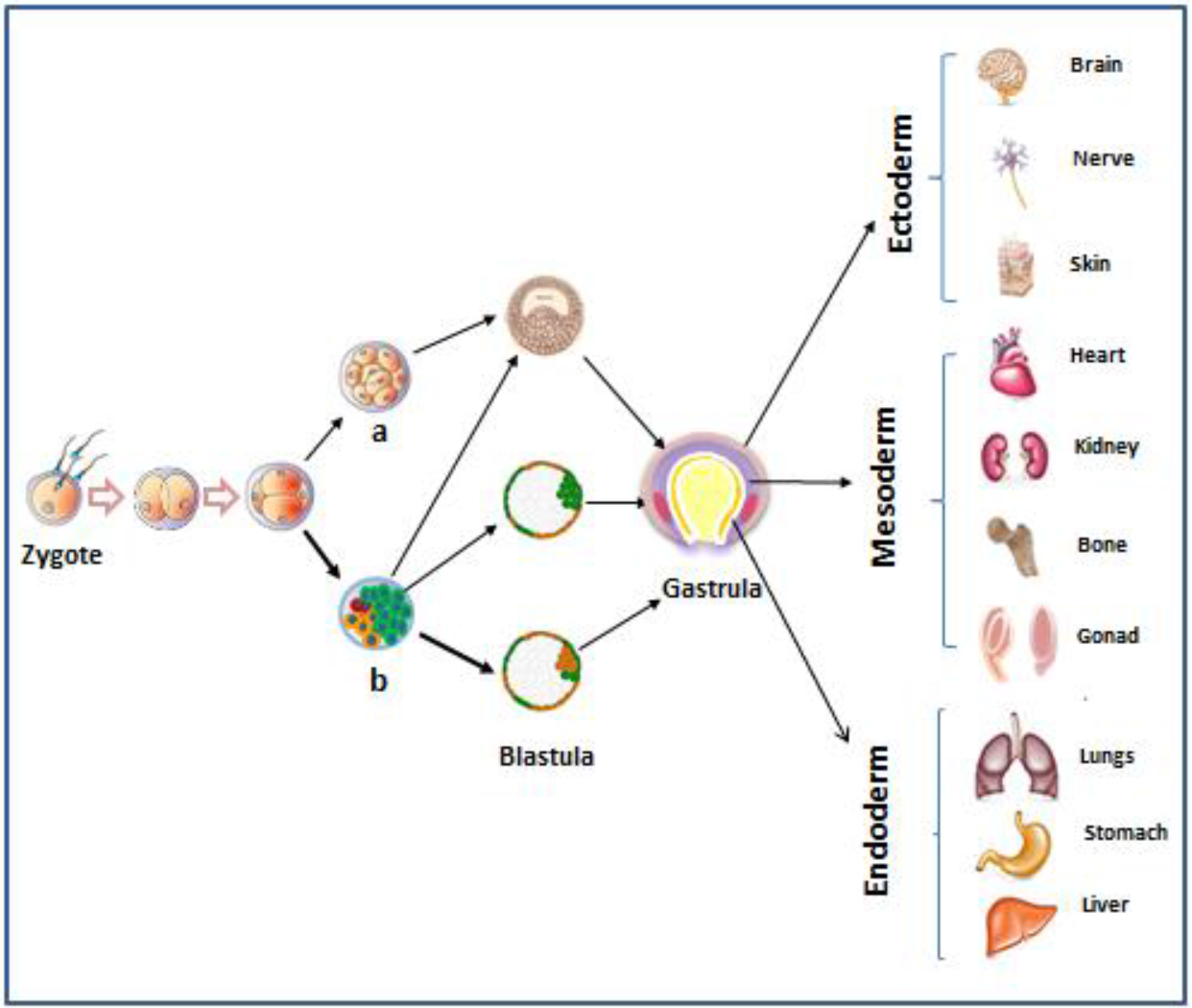
A simplified view of organogenesis. Zygotes with normal (a) cells and mutant cells (b). These could produce organs with normal tissues or mosaics (sectorial). Although various tissues are quasi-independent, they are connected by blood vessels and nerves. They possess modular and allometric properties (10), as well as show normal distribution, and behave as quantitative traits. It is assumed that sperm and egg cells carry a few ancestral mutations, and recruitments or abatement of cells follow demographic principles (see the text).

## Features of frailty

In the life course of individuals (i.e., from fertilization to death) and populations, senescence is primarily restricted to post reproductive stages, during which, senescent cells accumulate in multiple tissues and organs leading to vulnerability and pathogenesis of many diseases of somatic origin (21). Consequently, post-reproductive individuals start showing both physiological and functional decline, collectively termed frailty. From a gerontological perspective, frailty is a stochastic and dynamic process of deficit accumulation which occur ubiquitously at subcellular levels, ultimately affecting tissues, organs, and integrated organ systems, especially under stress. Although some individuals have greater propensity to accumulate deficits, on average, deficit accumulation varies across the life course and likely is mutable. (22). Since, cells are the primary sites of deficit accumulation, cellular frailty is a major cause of declining physiological and physical functions of organs (“organ deterioration”(23), resulting in single or co-occurrence of several late-onset diseases (multiple morbidity). Together, these processes contribute to reduced survival ability of such phenotypes (24–26). Thus, senescence, frailty and morbidities may share common origins (27).

## Mutations as catalysts of senescence

Among the many theories posited to explain the aging process or senescence, accumulation and amplification of inherited (constitutional) and acquired mutations in somatic and reproductive tissues are widely recognized as a cause of aging [mutation accumulation theory (MA) (28, 29)– (Medawar 1952; Szilard 1959). In a similar vein, Muller (30) suggested that adverse effects of numerous mutations in the genome, may cumulatively create a “load” (mutational load - ML) which could become a “burden” on the health of individuals in populations. The load is generated by sequential cell division initiated following post-zygotic fissions, which in turn create opportunities for DNA replication errors during strand segregation at every cell division event, some of which are repaired by DNA repair mechanisms. Nonetheless, the “burden” of somatic mutations also increases linearly with age. Although the burden of mutational load “could be measured only abstractly in terms of fitness, but … felt in terms of death, sterility, illness, pain and frustration” (31). Indeed, a few of these factors mediate both frailty and morbidity. Subtle differences aside, both MA and ML theories address the same issue; i.e., adverse effects of mutations on somatic and reproductive tissues, and ultimately, overall health.

The MA process is a gradual and ubiquitous feature among all differentiating cell lineages within tissues and organs, and organ systems within the organism – a process comparable to asexually or clonally reproducing organisms. In fact, “the dividing zygote is a clone. Just like a clone of unicellular organisms, embryonic cell divisions pass on any genetic variation which arises in previous divisions” (14). Each additional mutation may have cumulative and multiplicative adverse or advantageous effects on the organism at various levels: genomic instability, molecular heterogeneity, loss of cell-division, altered gene expression, impaired inter-cellular communication, tissue disintegration, organ dysfunction and vulnerability to stresses and all of which may synergistically influence both fitness and healthspan (32–37).

## Mutation accumulation in somatic tissues as a ratchet

Mutations appear to accumulate at a rate of 1-2 per division during early embryogenesis, and a cascade of additional mutations would be accrued in subsequent embryonic lineages (38) and ultimately develop into a complex organism. Consider the fact that an adult human is composed of 37 trillion cells distributed among approximately 80 organs (13, 39), and all of which connect back to once cell – the zygote. Correspondingly, differentiation and development of the zygote over decades would create ample opportunities for billions of mutations to accumulate virtually in every tissue and organs, despite DNA repair mechanisms. Like asexually propagated organisms, various classes of mutations [lethal, semi-lethal, nearly neutral mutations - (40, 41)], with wide-ranging and largely irreversible allelic effects gradually accumulate virtually among all somatic cells, cell lineages and tissues (42) in a stepwise ratchet-like manner(Figure 1, 2)., also known as Muller’s ratchet (43). Although, in principle, human cell lineages cannot readily produce a new organism, “inheritance and replication of (cells) is not by no means precluded” (14). The ratchet typically involves, “… an instability in the distribution of the numbers of carriers of deleterious mutations genomic instability when there is no recombination, brought about by genetic drift in a finite population” (44). Clearly, the drift of a finite number of cell population containing unstable genomes, among dividing population of cells within tissues and organs (comparable to chimeras) could lead to their diminished or complete physiological dysfunction/shut down, affecting the viability fitness of cells and their lineages – a process analogous to meltdown or extinction (45). Harmful effects of mutation accumulation might be amplified further under stressful conditions (46) such as internal and external environmental insults. The inter-cellular spaces left by dysfunction and death of cells, may remain unoccupied by newer cells or may be filled with toxic cellular waste products. Together, these processes could potentially reduce intercellular integrity and robustness of the cell and tissue architecture and interfere nutrient signaling and distribution, initiating senescence (47, 48). At this stage, cells could enter into an irreversible physiological state, and can no longer divide. Senescent cells, on the other hand, also secrete various proteins such as inflammatory cytokines and growth factors, which could stimulate the resident stem cells to repopulate and remodel the tissue (25). Thus, senescence could be both a destructive and a constructive process. In older tissues, however, it is largely a degenerative process.

From a genotype-epigenetic – phenotype map (17, 18) perspective, it is reasonable to suggest that lifelong, cumulative and differential effects of the accumulation of ‘deficits’ at the genetic, epigenetic and cellular phenotypic levels affect the physiological and genetic homeostatic properties of cellular phenotype and ultimately the aging phenotype and phenotypic components (49–51), accordingly. Upon superimposing the concept of frailty onto the principle of Muller’s ratchet we postulate that relative frailty of phenotypes [(i.e., cell(s)], tissue and multimorbidity of organs within individuals) may be a function of the load of mutations they carry. Under nearly neutral theory, both synonymous and non-synonymous mutations impart a different degrees of deleterious effects (in the absence of advantageous back mutations) on the phenotype (34, 52). Then, in a demographic sense, the degree of frailty of tissues, and organs may reflect a range of susceptibilities to mutational burden they carry (30, 43) in relation to stage and age of tissues. Ultimately, organ failure may represent a functional collapse at a “tipping point”(24) leading to physiological and dysfunction of tissues – simply, “mutation meltdown” (45).

It has been suggested that understanding possible genetic and molecular mechanisms of the frailty process will be instrumental in order to develop preventive and palliate measures to counter certain aspects of morbidity (53) in multiple medical specialties (54). Clearly, ideas from evolutionary genetics could be extended to gain insights on some of the enduring questions in aging and human health. Indeed, “Some patterns in metazoan developmental processes may be approached by quite standard neo-Darwinian cognitive styles.”(14). To our knowledge, a comprehensive study of evolutionary genetic bases of the relationship among cellular senescence, frailty and multimorbidity has not received much attention. We have two objectives in this study: a) to apply Muller’s ratchet principle to aging process in somatic tissues, and b) to discuss the utility of evolutionary genetic mechanisms toward understanding the origins of frailty, morbidity and senescence. We make the following assumptions: 1) cell is the primary unit of differentiation, development, and regeneration and also represents correct abstraction of the individual(55, 56); 2) All organs and mosaics within an individual are derived from the same zygote (cell), yet they are differentiated, connected, function autonomously, and might be exposed to the same or different environmental conditions; and 3) cellular senescence (57) is both a norm and a precursor to other known degenerative processes in tissues, organs and ultimately the organism; but could also influence regeneration and tissue remodeling (25, 47). Among numerous theories advanced to explain the aging process (58), we focus only on Muller’s ratchet as a possible cause of frailty, vulnerabilities to diseases, and morbidity in the course of human growth and development.

## Model

Our model takes a demographic approach to aging process starting from a single stem cell through differentiation and development into mature tissues and organs, analogous to embryogenesis and its clonal lineages (14), as well as their gradual loss of physiological functions over time, and eventual death (Figure 1). It assumes that each cell is specified by three parameters, the rate of cell division, functionality or healthiness and death rate (respectively, denoted by *b*, *f,* and *d*), which are given by a function of the mutations that the cell has accumulated. Basically, by assuming all mutations exert deleterious effects on cell populations to varying degrees (40, 41), *f* and *b* decrease and *d* increases as a cell accumulates mutations. *f*_0_, *b*_0_, and *d*_0_ represent their values at birth, and their changes by a mutation is given by *δf*, *δ*b and *δd* (see Suppl. for details). *F(t)*, *B*(*t*) and *D*(*t*) are the sums of *f*, *b* and *d*, respectively, over all cells in the population at time *t*.

As illustrated in Figure 2, any tissue is a population of cells, in which each of the constituent cells are genetically variable (hence different genotypes) and some are more variable than others. The population proliferate from a single cell, and the cell population size at time *t* is denoted by *N*(*t*), and *t* represents age and *t* starts at *t* =−1 (roughly corresponding to the time of fertilization and zygote formation). *N*(*t*) is assumed to increase up to *K*, which represents the population size of an adult. Then, as aging proceeds, *N*(*t*) could start declining. This assumes well-known phases of density dependent population growth (lag, stationary, decline), as well as exhibit the salient features typical to such populations - connectivity, cooperation, competition and coevolution (59, 60). Thus, the process may be divided into three phases (Figure 2B). The first phase (Phase I - lag) is from the initial state to *T*_1_, after which *N*(*t*) = *K* (Phase II - stationary). *T*_2_ is the time when *N*(*t*) starts declining, and this phase is referred to as Phase III.

**Figure 2.**
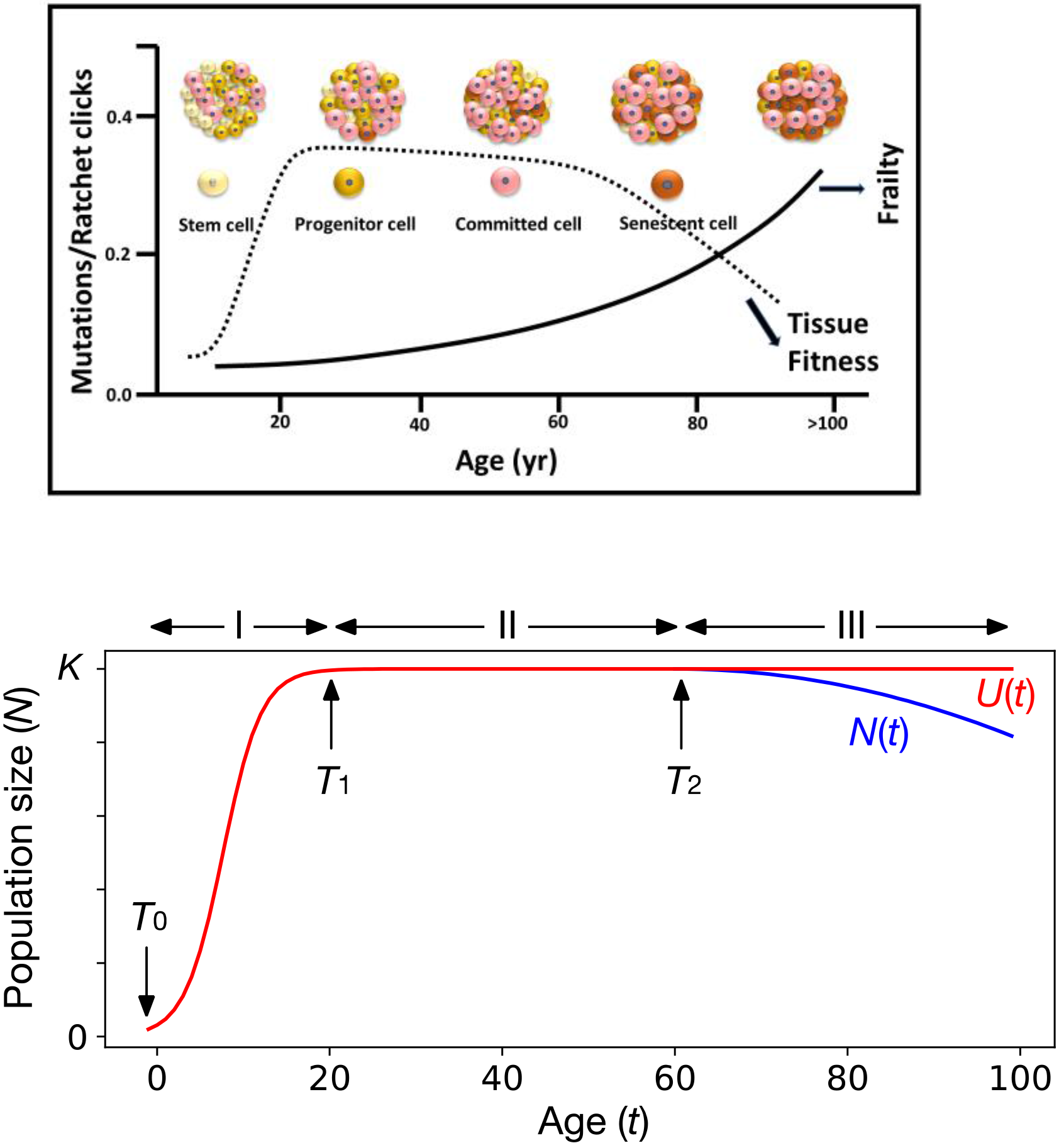
**A:**Some general features of mutation accumulation as ratchet, cellular senescence and frailty among somatic tissues in relation to age: hematopoietic cells (104), incidence of cancer, multimorbidity caused by cardiovascular, cognitive, respiratory disorders (105–107), multimorbidity and the frailty index (108). Cellular diversity within tissues modified from Horvath 2018. ***Muller’s ratchet clicks more with age. Also, the fitness of cell populations is rank-ordered and the average excess of deleterious mutational effects (Crow 1986) are greater in a given tissue with advanced age, which is equated with frailty‥* B:** Our model of the cell population size, which introduces *U*, the upper limit of the population size (red line). *K* is the adult population size, so that the population size (red line) increases up to *K* in Phase I (*t* < T_1_=~20 yr). Then, in the adult phase (Phase II), the population size is stable at *K*. When the aging process proceeds and it becomes difficult to maintain the population size at *K*, the population size decreases in Phase III (blue line), thereby accelerating the aging process.

In this work, we modified the Moran model such that we can incorporate the change of population size as illustrated in Figure 2B. In brief, we introduce a parameter *U*(*t*), an upper limit of the population size (e.g., all healthy tissues and organs have a species specific size limit). The population can increase up to *U*(*t*) if *B*(*t*) > *D*(*t*), but the population will decrease in size if *B*(*t*) < *D*(*t*). When the tissue is young and healthy enough (*B*(*t*) > *D*(*t*)), the population size is kept at U, but as all cells in a tissue accumulate deleterious mutations and *D* becomes greater than *B*, the population size starts declining. See supplementary part for theoretical details.

## Results

Our model incorporating demographic changes provides a simulation framework for exploring the aging process in growing cell populations. In addition, some analytical expressions on the nature of mutation accumulation during aging process may be obtained with simple and biologically justifiable assumptions. We first describe the theoretical results to gain insights into the mutation accumulation process through aging process or cellular senescence (4), followed by more detailed analyses by simulations.

### Theory

Let us first focus on Phase I, in which *B*(*t*)/*D*(*t*)>>1 and *D*(*t*) is so small that most events are cell divisions due to an increase of *U*(*t*) and *N*(*t*). We then derive approximate expressions for the number of somatic (*S*) mutations, *S*_*ave*_(*t*) and *S*_*total*_ (*t*), the average and total numbers of mutations that cells have accumulated by time *t* (i.e., *S*_*total*_(*t*)= *S*_*ave*_(*t*)*N*(*t*)). Consider a situation where there are *N* cells waiting for an increase of *N* to *N*+1 by division. If a cell having *s* mutations (constituent) undergoes a cell division, its two daughter cells will have *s* mutations plus new mutations that occur at rate *αμ* in one cell and at a rate (1-*α*)*μ* in the other.

Therefore, when *N* increases by one through a single cell division event, the expected increase of *S*_*total*_(*t*) is *S*_*total*_(*t*)/*N*(*t*)+μ. Thus, *S* increases as *N* increases, and if we ignore death events, *S*_tota1_(*t*) roughly satisfies

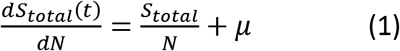

with a continuous approximation for a large N, from which we have

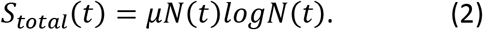

*S*_*total*_(*t*) is thus given by a function of *it* if *N* (*t*) (or *U* (*t*)) is specified (e.g., see equation (S1 in supplementary information).

If cell death (apoptosis and necroptosis) is taken into account, we consider two paths from *N* to *N*+1 by assuming the time interval (*Δt*) from *N* to *N*+1 is so small (i.e. time taken for one cell division) that at most one cell death occurs in *Δt*. The first path is that a single cell division increases *N* to *N*+1, as described above. The second path involves a cell death, and the dead cell is eliminated by intercellular competition, which changes *N* to *N*−1. Then, two cell divisions follow and the population size becomes *N*+1, according to the assumption of a very small *Δt* (so that multiple death events are not considered). Let *p* be the probability that the system goes through the second path. Because the expected increase of mutations is *S*_*total*_(*t*)/*N*+2*μ* in the second path, Equation (5) may be written as

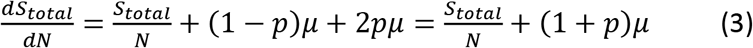

where, *p*=*D*(*t*)*Δt*. *Δt* depends on the growth function, *U*(*t*).

The process of mutation accumulation in phase II is quite simple because a death event is always coupled with a cell division. That is, the accumulation of mutations simply depends of the death rate, *D*(*t*):

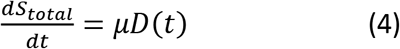

This process is analogous to mutation accumulation in constant-size asexual populations also called, Muller′s ratchet, which reduces fitness (61).

In Phase III, a dead cell may not be filled by a new cell, which may have a cascading effect on the integrity of cell populations: formation of intercellular spaces, disintegration and interference with their adhesion, connections and communications (62) as well as deceleration of the repair process. Let this probability be q= *B*(*t*)/*D*(*t*). Because such an event simply decreases a cell,

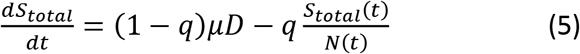

It is quite straightforward to compute *S*_*ave*_(*t*) from equations (1-5) if the effect of mutations on *b*, *d*, and *f* is ignored (i.e., mutations are neutral). Under this setting, cells do not degrade such that Phase III does not arise. With this assumption, Figure 3 demonstrates that the prediction of equations (1-5) is in excellent agreement with simulation results detailed below, indicating our theory well describes the accumulation of mutations in Phases I and II. Simulations were performed by assuming a logistic growth of the population size up to *K*=100 with the initial growth rate *r*=0.5 [(equation (S1) in supplementary information)]. We assumed *b*=0.5, and five values of *d* were considered (*d*=0, 0.01, 0.02, 0.05, 0.1, from bottom to top in Figure 3). In all five cases, the number of mutations, *S*_*ave*_(*t*) dramatically increases in Phase I. In Phase II, *S*_*ave*_(*t*) linearly increases and the speed of ratchet depends on the death rate; *S*_*ave*_(*t*) increases faster for a larger *d*. This is simply because a cell death is always followed by a cell division, through which mutations are introduced (47). Therefore, a large *d* implies a fast turnover of cells, resulting in a fast accumulation of mutations. Our theoretical result for Phases I and II indicates that, (i) when the tissue grows to maximum (adult) size, a large number of cell divisions occur and mutations accumulate very rapidly, potentially damaging the functionality or healthiness of the tissue. (ii) The rate of mutation accumulation totally depends on the rate of turnover (death rate) when the tissue sizes reach the adult size. Therefore, for tissues like brain where the number of cells are almost fixed (relative to other organs such as liver, musculoskeletal system) and little of no cell division occurs in Phases II and III, mutations in very early stages in Phase I plays a crucial role, particularly in the aging of the brain. This result is consistent with a recent finding that most somatic mutations in brain originate first several rounds of cell divisions. In contrast, tissues with a high rate of cell renewal (e.g., skin, blood, small intestine - see above), the relative contribution of mutations at such very early stages in Phase I is generally small because the tissue continues to accumulate mutations throughout the lifespan of individuals.

**Figure 3.**
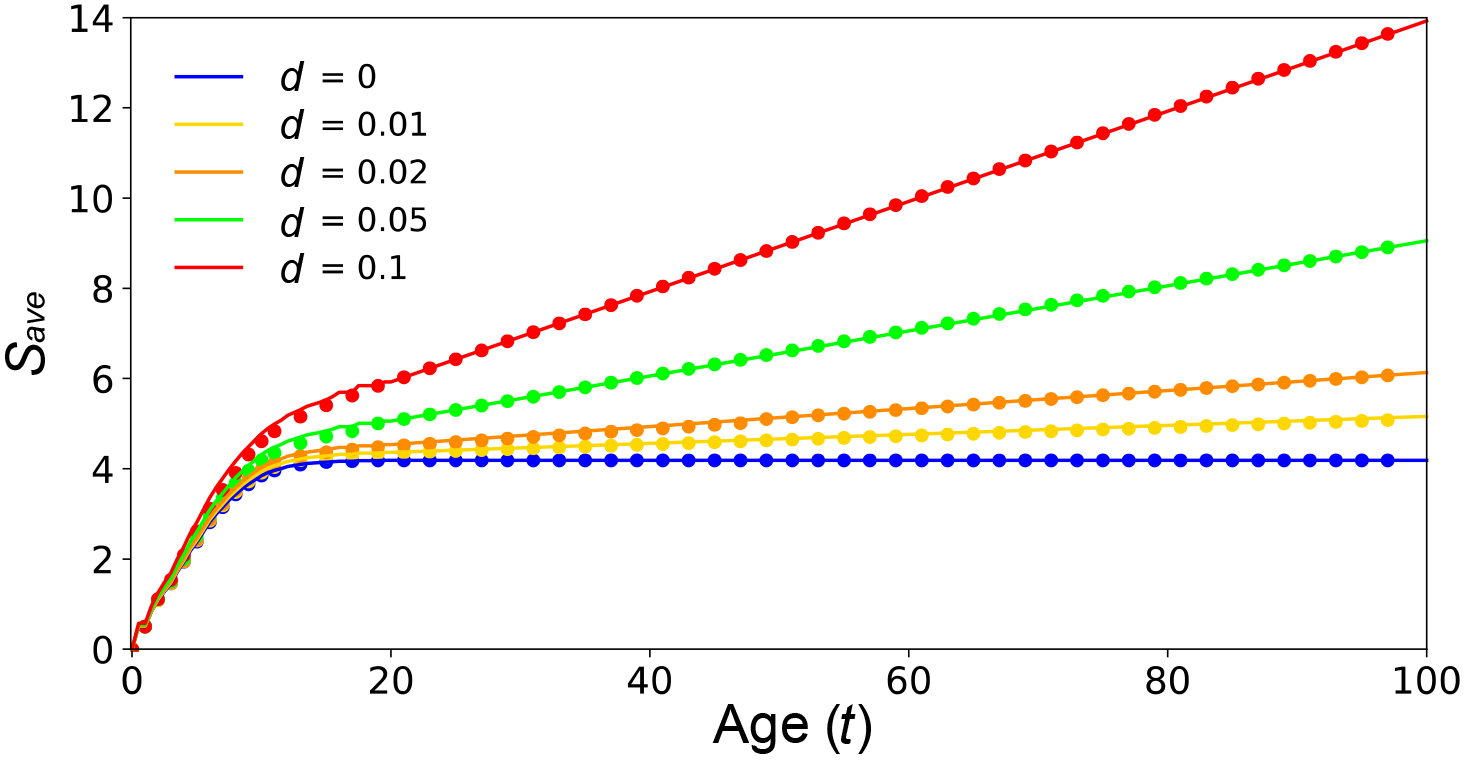
The number of accumulated mutations per cell, *S*_*total/n*_ = *S*_*ave*_ (y-axis) as a function of T, age. Theoretical prediction from equations (1-5) is shown by solid lines and simulation results are shown by closed circles. A logistic growth of the population size up to *K*=100 with the initial growth rate *r*=0.5 is assumed (See equation (S1) in Supplementary Information). We assumed *b*=0.5, and five values of *d* were considered (*d*=0,0.01, 0.02, 0.05, 0.1).

### Simulation

The above theory is based on the simplified assumption that mutations do not affect *b*, *d*, or *f*, therefore it is applicable when the tissue is healthy (homeostatic state) enough to maintain the cell (approximately constant) population size (i.e., *B>D*). In order to explore the entire aging process, we performed simulations under a more realistic situation, where the effect of mutations on *b*, *d* and *f* is involved. With this effect, *f* and *b* decrease but *d* increases along with the accumulation of mutations as we defined in equations (1-5). As a consequence, *D* eventually becomes larger than *B*, and from this time point, the phase can shift from Phase II to Phase III. Because it is not very straightforward to evaluate *S*_*ave*_(*t*) with the above equations, we performed a number of runs of simulations. As well as Figure 2, *r*=0.5 and *K*=100 were assumed. For convenience, it is assumed that a deleterious mutation affects *f*, *b* and *d* equally, that is, *δf* = *δb* = *δd* = *δ* = 0.05. *b*_o_=0.1 was assumed and we changed *d*_0_ to vary the speed of aging.

The consensus we found in a number of simulation runs is that the tissue degrades gradually over time, as expected for frailty, and that it will eventually reach physiologically dysfunctional state [(meltdown, sensu (45)] both by the shrinking population size of healthy cells and with the death of apoptotic and necroptotic cells. In our model, incipient meltdown could begin when the existing cells do not have ability to fill all empty slots, that is, when *D*(*t*) exceeds *B*(*t*) (the point at which the blue and red lines cross in Figure 3). After this time point (Phase III), the population size decreases gradually and irreversibly. In this phase, the aging process accelerates largely because, in addition to the decrease of healthiness of each cell, the healthiness (^~^fitness) of cell population decreases synergistically together with the decrease of the population size. It is also interesting to note that there is great heterogeneity in fitness (or vulnerability) among cells within the cell population. That is, in principle, cell populations diverge into clusters due to mutation accumulation within a tissue or among tissues nested within an organ, and could be graded in relation to their viability fitness (63). Cell is the driving force for the ratchet to proceed. A low-ranked cell and cell-lineages likely become physiologically dysfunctional and die, and in a healthy tissue (Phases I and II), a daughter cell of a high-ranked cell likely replaces, but cell division of a high-ranked cell (due to high mutational load; see Figure 2) would decrease the fitness of the two daughter cells and their lineages. Such deterioration of the fittest cell signifies a “click” of the ratchet, and the deterioration of cell populations is proportional to number of clicks (64). The situation is even worse in Phase III, where high-ranked cells are too deteriorated to replace low-ranked cells, thereby causing a decrease of the population size (therefore the tissue healthiness dramatically drops; Figure 2).

## Discussion

Physiological and morphological decline with advanced age is universal to all organisms. Some of these deteriorative processes, commonly described as frailty, are apparent at multiple levels of biological organization ranging from sub-cellular to supra-cellular systems: genome, epigenome, tissues, organs and the organism. Clinical frailty involves a cumulative decline in the developmental, anatomical, physiological and phenotypic functions and numerous human disorders (25). It is attributed to “lifelong accumulation of molecular and cellular damage”(49, 65, 66) which could disrupt genomic and biochemical networks leading to dysfunction of a cell or a tissue, an organ or many. Clearly, although frailty is recognized primarily as a geriatric condition it is an emergent condition and a syndrome affecting multisystems of the human body at various biological levels and stages in the entire lifespan of individuals (67). Damage at the molecular level includes both benign and pathogenic variants among arrays of inherited (constitutional) and somatic (passenger) mutations (68) in the genome. These variants accumulate differently in the genome among the diverging lineages of cell populations in relation to age - a process analogous to mutation accumulation in asexual lineages (69). In fact, cumulative effect of constitutional and passenger mutation could create dosage imbalances and epigenetic drift, which in turn, are known influence organs and organ systems in aging, cancer and other clinical conditions in hereditary diseases (36, 70).

In the present study, we extended the Muller′s ratchet principle on growing and branching population of cells assuming that they live for the entire life course (say 80 – 100 years in humans) in several steps: First, Moran birth-death (demographic) process of cells in finite cell populations (i.e., tissues); second, we introduced mutations into these cells at various stages of development of individual cells and simulated the consequences of these branching clonal lineages of cell populations. Since mutations accumulate differentially both within and among cell populations (71), with time, they become structured. Accordingly, it is reasonable to assume each individual cell population could show different viability fitness, however small it may be. Our results agree with empirical studies dealing with mutation accumulation and trajectories of senescence on model systems as well as human disorders. First, growth, stability and decline of cell populations follow features typical to stable populations with constant age structure. This pattern also agrees with age-dependent gene expression profiles, which often serves as a proximate index of genomic variation (12, 32). Second, as shown in the bottom panel of figure 4, birth (blue line) and death rate (red line) increase or decrease in relation to mutation accumulation. These lines cross around age 50 [(essential lifespan, (37)] after which the cell populations enter a declining phase (*i.e., B<D*). This trend also corroborates with the Hayflick limit, which suggests that normal healthy cells in culture will divide between 40 and 60 times before reaching a plateau afterward their density declines due to senescence. Third, if cells are dividing abnormally fast (Fig. S1C), they might survive for about 15 years (e.g., Hutchinson-Gilford syndrome). However, it is likely that the health status of various tissues and organs could vary, in relation to relative degrees of mutation accumulation. Interestingly, this is generally the case for progeroid syndromes, in which a causal relationship with gene specific accelerated rate of mutation accumulation and attendant aging syndrome have been established. In these syndromes, mutations accumulate differentially and show pleiotropic effects among tissues and organs affecting many physiological pathways, frailty and ultimately affect their health and longevity(72). Alternatively, our simulation study also showed the possibility for renewal or a capacity for homeostasis, suggesting that while senescent cells die, dead cells may be replaced by new stem cells opening up an opportunity for rejuvenation or remodeling of the tissue. How might these two seemingly opposite phenomena co-occur? Note that when senescent cells are eliminated from a tissue, senescence associated secretory phenotype (SASP) recruits T helper lymphocytes. The SASP remobilizes and repopulates the nearby progenitor cells and eventually remodel the tissue (25). Clearly, under special conditions, senescence is both a degenerative and creative process and help restore damaged tissue (25, 73) and not a consistently unidirectional process. Four, mutations that originate first rounds of cell divisions appear to have greater influence of cellular senescence than the ones that appear late in the lifespan of the tissue or organ. Recent work suggests that mutations that occur prenatally in the brain could amplify later into life into mosaics and cause neurodegeneration (74, 75) Interestingly, mutations rate as measured by single nucleotide variants (SNVs) differed vastly between the mutated cell line (10^−4^) vs. 10^2^ – 10^3^) in normal neighboring cells. This result not only supports the principle of Muller’s ratchet, but also the origin of latent and aggressive chronic disorders, as well as shifts in the rate of mutation accumulation in relation to age and stage in the development of the organ. This may also explain about four-fold increase in multimorbidity after 65 years of age (65) relative to younger ages.

**Figure 4:**
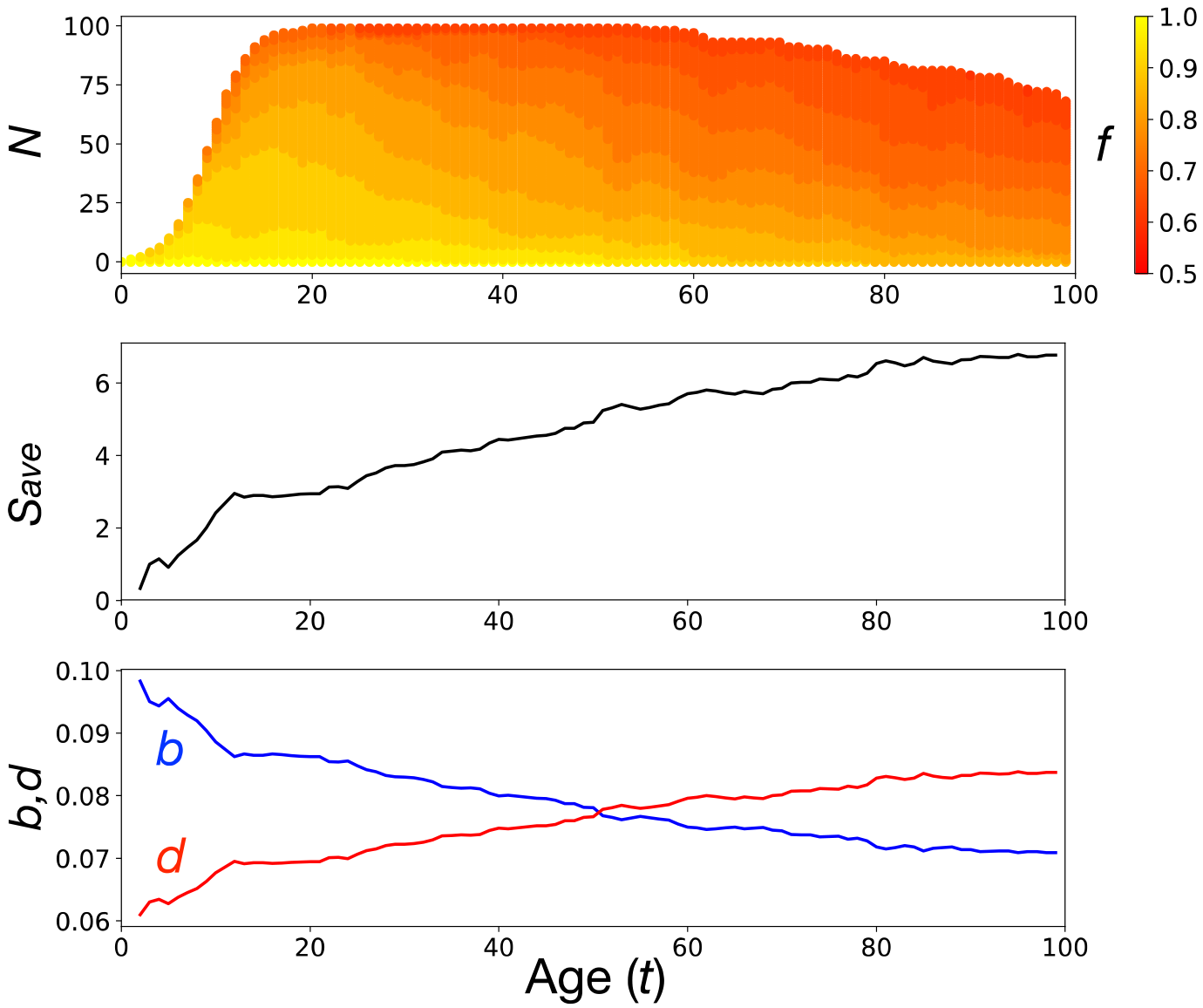
A typical simulation run. *d*_0_=0.06 is assumed as the initial condition. As *B*(*t*) decreases and *D*(*t*) increases (bottom panel) with accumulating mutations (middle), and they became nearly identical when cells accumulated roughly seven mutations around *T*=50. From this time point, the population size can start shrinking. Along the run, cells degrade with decreasing functionality (*f*), from yellow to red in the top panel

As a parsimonious explanation, Muller′s ratchet among cell populations, illustrates the possibility that specific cell populations within tissues and organs could accumulate numerically excessive individual mutations regularly and genetically diverge over time. Hence, such an ensemble of cell populations would share similarities with structured biological populations, with the only exception that cell populations may or may not experience the amount or spectrum of environmental stresses that the latter ones do. The collective ensemble of the deleterious influence of all mutations in the genome (not necessarily only the most harmful ones) in genetically distinct somatic cell lineages and populations may also contribute to reduced viability fitness specific to each population. Further, these populations may also display different intrinsic vulnerabilities for degeneration, within a given tissue or organ, due to different levels of genetic variation among them. Many of such early events that occur during the process of mutations load in small cell populations could be common to both senescence and cancer (76). This situation is comparable to Fisher′s ″average excess″ (77) defined as the amount by which the weighted fitness of individuals carrying specific alleles exceeds that of the population as a whole (78). The only difference is that, in our case, we have populations of individual cells, instead of mature individuals in populations. Note that the average excess represents an average of allelic products (proteins/metabolites) a cell or a cell population produces; which could increase or decrease its fitness depending upon the phenotypic effects of mutant alleles carried by each individual (78, 79) (Figure 2). This scenario also hints at the possibility that in some situations, total mutation rate, and not the deleterious effect of individual mutations may influence the average fitness of cell populations as illustrated by the Haldane-Muller principle (80). These spectrum of mutations could potentially induce various degrees of cis-ruption in the spatially organized network of genes and haplotype networks which could progressively retard or even shutdown cellular repair mechanisms (81), and metabolic flux (36). These synergistic and time-lag processes might also exert cascading and disintegrating effects on subtle connections, cooperation or competition among intra - and inter-cellular organization, communication and transport; which may result in the genetic divergence of somatic tissues (82) and subsequently influence the frailty among physiological, anatomical and morphological traits. In fact, while networks confer stability to biological systems (83), instability could lead to multiple organ dysfunctional syndrome (MODS (67)]. Obviously fitness could change among the divergent population (crypts) of cells, within tissues in relation to number of mutations (31, 83). An excellent example of this process was recently provided by (84)(85), epilepsy (86), cardiovascular and skeletal disorders (21) as well as several brain disorders including cerebral palsies (87) and cortical malformation (88). A comprehensive list of diseases associated with many organs and their component tissues (25) clearly suggests phenotypic manifestation of interaction among developmental disorders and frailty leading to multimorbidity. A more recent study involving more than 3000 families (89) and coworkers have reported that diseases of de novo origin may be more common than previously thought, and could affect nearly four million children worldwide, annually. Similarly, somatic origins of late onset complex disorders such as stroke, demyelination, optic nerve disorders and brain cancers represent only the “tip of the iceberg” (90) of numerous disorders, yet to be described.

There are limitations to our study, however. Since our goal was to extend Muller’s ratchet principle to link cellular senescence and frailty, we modeled these processes, based on average mutation accumulation in populations of brain cells as example. The model could be extended, however, to other tissues assigning tissue specific mutation rates (91), and recruitment cycle of cells, which are often context dependent, as discussed. We have relied on mutation accumulation process in asexual lineages, in which Muller′s ratchet has been suggested to operate. In our model, the growth parameter for cells, ″*U*″ is greater at the initial stages (phase 1 – childhood), later it becomes fairly stable, but decreases at later ages. These results agree with molecular data (38, 75) as well demographic and evolutionary models of aging (6, 92, 93). For instance, Lodato et. al (75) reported approximately 1000 SNVs by 5 months, but steadily increase after 40, and exponentially so after 80. They call the plateau ″quiescent phase.″ Note, the quiescent phase envelops region of maximum Darwinian fitness (6). Additionally, it is important to experimentally determine fitness differentials among cell lineages within tissues in relation to mutation accumulation (83).

Another limitation is that we considered a population of stem cells, which accumulate mutations when a stem cell undergoes cell division to produce another daughter stem cell. However, there are at least two factors that could induce mutations into stem cells. One is mutations that arise during the division and differentiation of stem cells, and the other is mutations that are induced by the micro-environment of the tissue (94). These two factors can be incorporated in our derivation (equations 1-5) by replacing by *μ* and *μ’,* where *μ’* is defined as the total mutation rate including these two factors. These two factors well explain the observed exponential increase of SNVs after 80 yr. in brain (75). They also explain the recently reported almost star-shaped tree of hematopoietic stem cells (38). The tree indicates that the hematopoietic stem cell population grows rapidly and reaches a plateau, and since then they accumulate mutations by producing additional progenitor cells.

In conclusion, the Muller′s ratchet principle provides a rational foundation to study the origin, maintenance or augmentation of vulnerability, frailty, morbidity and mortality associated with aging as well as many inherited and latent disorders. The principle focuses on generally predictable effects of mutation accumulation on the aging phenotypes and compliments two well-known mechanisms of aging – the Hayflick limit and mutation rate (91, 95). This trend is also reflected at the genomic level in hematopoietic cells (38). Note that mutation rates, telomere shortening and the Hayflick process represent, respectively, mutational, molecular, and cellular clocks. Similarly, aggregate methylation status shows a linear relationship with age, also called, DNA methylation (epigenetic) clock (8). Mutational, molecular, cellular and epigenetic clocks are largely invariant and species specific “clock-like” processes among healthy organs and individuals (96). Similarly, at the organism level, gestation period, childhood, puberty, ovulation and menopause are predictable traits. Note that puberty, menopause and life span are evolutionarily conserved landmarks in human life history (97). It is not surprising that methylation, which links the G-P spaces has been shown to predict biological aging and longevity (8, 32). From a G-E-P map perspective, these are suggestive of the predictability that exists among levels of biological diversity (16). On the other hand, epigenetic drift caused by stochastic changes in methylation in aging cell populations (36) may reflect the underlying genetic drift of those populations due to mutation accumulation. Epigenetic drift could cause departures from this norm, however, by many orders of magnitude, among tissues, organs and stress (8, 74, 98) and disease conditions.. Therefore, in principle, there may be a predicable relationship between the pace of “clicks” and the “ticks” of Muller’s ratchet and epigenetic clocks. Deceleration or acceleration of these regular events could serve as indices of health, morbidity and senescence.

How might this information be used in a clinical setting? It has become a routine practice to classify mutations into finite number of classes such as pathogenic, benign, uncertain and actionable. Instead, in some cases, and in accordance with the Haldane-Muller principle, cumulative effect of the entire spectrum of mutations may have to be considered in relation to demographic and developmental aspects of cell populations/tissue, for developing tissue level interventions within organs. Accordingly, guidelines already developed for clinical diagnostic purposes based solely on the effect of individual mutations, needs to be reevaluated. Here aging and cancer related vulnerabilities would share both similarities and differences (99). Perhaps, suppression or elimination of the effects of the entire spectrum of mutations by novel means might be helpful toward ameliorating overall health (100). Further, detailed investigations into the origins, distribution and effects of somatic mutations could open up the possibility for describing many medically unexplained symptoms and even new disorders on the basis of novel mutations and the degree of mutation load. These might also require developing novel clinical trials in order to manage medically undiagnosed symptoms at those levels of diversity. In other words, the scope of N-of-One clinical trials (101) could be extended to include interventions at any level of biological hierarchies within an individual. For instance, from the perspective of precision medicine, it holds promise in order to effectively identify individuals at higher risk of developing singular or multiple morbidities (102) and for devising new preventive measures such as tissue and organ specific synolytics (3, 103). Clearly, some concepts of evolutionary genetics have the potential to illuminate enduring questions in human health and longevity.

## ACKNOWLEDGEMENTS

We dedicate this paper to Richard Lewontin and Tomoko Ohta and for their prescient ideas on units of selection and mutational effects, respectively. DRG thanks Peter Ellison, Nir Barzilai, Jan Vijg for support, and Dr. Sri Raj for suggestions.

## Supplementary information

### Assumptions and Parameters

Our model takes a demographic approach to aging process starting from a single stem cell through the development of a mature tissue, and its reduced loss of physiological functions over time, and eventual death. We model the aging process such that it is assumed that a cell should be specified by three parameters, the rate of cell division, functionality or healthiness and death rate (respectively, denoted by *b*, *f*, and *d*), which are given by a function of the mutations that the cell has accumulated.

Let *b*_0_, *d*_0_ and *f*_0_ be these parameters at birth. Then, these parameters for an individual cell with *k* mutations are given as follows. Assuming that mutations make the cell unhealthy, f is given by

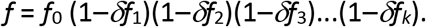

where *δf*_*i*_ is the relative reduction in the functionality caused by the *i* th mutation. Similar should be applied to the cell division and death rates. As we assume mutations cause a decrease of the birth rate and an increase of the death rate, we have

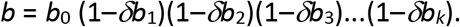

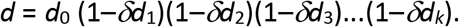

These equations mean that a single mutation is specified with its effects on the three parameters.

We here assume that mutations occur during cell division, that is, a cell lineage splits into two, and they accumulate mutations due to replication errors and random factors at rates *αμ* and 846 (1-*α*)*μ*. To be simple, two types of mutation could be involved, lethal and deleterious, and the proportion of lethal mutations is denoted by β. The former is an extremely damaging mutation, with which the cell will die and eliminated by inter-cellular competition (109) or apoptosis. The latter is a deleterious (or neutral) mutation that affects the cell viability (^~^ cellular fitness) only partially (110), so that the cell becomes slightly less fit relative to its neighboring cells. The accumulation of deleterious mutations also causes a cell death in two ways. One is a random death according to its death rate (*d*), and the other is a forced death. The latter occurs when the cell has accumulated too many mutations so that the parameter of healthiness (*f*) becomes lower than a threshold value (*Lf*). *F*(*t*), *B*(*t*) and *D*(*t*) are the sums of *f*, *b* and *d*, respectively, over all cells in the population at time *t*. Under this setting, we model the aging process of a tissue along the accumulation of deleterious mutations in the cell population as detailed in the next section.

### Modeling a cell population

Our modeling is based on the Moran model, which assumes that an individual cell dies and then the empty slot is filled by a new cell. In other words, death and birth events are equal, so that the population size is kept constant. Cell death occurs at a constant interval. In this work, in order to incorporate the changes of population size, we add two modifications to the Moran model. First, cell death occurs randomly at any time point, depending on the death rate of each cell (not in a constant time interval). A dead cell may (or may not) be replaced by a daughter cell of one of the cells in the population, and the parental cell is randomly chosen according to its relative contribution to the population birth rate, *B*(*t*). Second, in addition to cell death, our model creates empty slots when *U*(*t*) increases. When an empty slot arises at time *t*, it is assumed that the slot is immediately filled (as the Moran model does) when the population has sufficient reproductive ability, that is, *B*(*t*)/*D*(*t*)>1, otherwise the probability of successful reproduction is given by *B*(*t*)/*D*(*t*). For simulating the aging process, the system starts with *B*(*t*) much higher than *D*(*t*), such that empty slots are immediately filled in Phases I and II. In our setting, *B*(*t*) and *D*(*t*) monotonically increases and decreases as the process proceeds, so that *T*_2_ is considered as the time when *B*(*t*) first becomes lower than *D*(*t*), (i.e., *B*(T_2_) ~ *D*(*T*_2_)). Thus, we cannot explicitly specify when the phase shifts from II to III. Rather, Phases II and III are defined such that phase III starts when *N*(*t*) starts declining. Therefore, *T*_2_ could differ among independent realizations with the identical parameter set. In Phase III, as a new cell can arise only when a cell dies in Phase III (as well as Phase II), once a slot is not successfully filled, it remains empty for the rest of the process, thereby resulting in an irreversible decrease of the population size (e.g., osteoporosis, sarcopenia, neurodegeneration in Alzheimer’s diseases). The process is terminated when the cell population (tissue) reaches functional attenuation due to dysregulation of physiological processes (melts down sensu (45) Lynch et. al 1993), that is, *F*(*t*) first hits a threshold value, *LF.*

assume a logistic function for *U*(*t*):

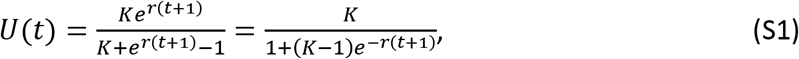

where r is the initial growth rate (cell division rate per year) and K is carrying capacity that roughly represents the population size of the adult tissue. We will later discuss the effect of this arbitrary choice of logistic function. In phase II, *U*(*t*) = *N*(*t*) = *K*, while in Phase III, *U*(*t*) = *K* and *N*(*t*) < *K*.

